# Standardized and rapid protocol for drought tolerance screening at seedling stage in small grain cereals and their wild relatives

**DOI:** 10.64898/2026.09.14.751556

**Authors:** Santosh Gudi, Jatinder Singh, Upinder Gill, Rajeev Gupta

## Abstract

Drought stress severely threatens the stability of production of cereals worldwide. Improving genetic tolerance to drought stress requires detailed characterization of global germplasm collections and their wild relatives under drought conditions. However, the lack of rapid screening methods remains a major obstacle in achieving this goal. Here, we present a rapid, reliable, and highly reproducible soil-based screening protocol for evaluating large germplasm collections under drought stress at the seedling stage. This protocol efficiently discriminates drought tolerant and sensitive genotypes with minimal environmental variability and is broadly applicable across various small grain cereals, enabling the evaluation of 1200 genotypes of wheat, oat, barley, and their wild relatives such as *Aegilops tauschii* and *Ae. umbellulata*. Additionally, evaluation of two contrasting wheat genotypes identified using this protocol under drought stress imposed at anthesis and grain filling stages provided evidence that their contrasting drought responses were also consistently maintained at the later developmental stages. Given its efficiency, reproducibility, and cross-species applicability, this protocol offers a practical approach for preliminary screening of large germplasm collections as well as for identifying contrasting genotypes to be included in subsequent breeding, physiological, and genetic studies.

## 1. Introduction

Cereals, mainly cultivated as a major source of human food and animal feed, are the members of grass family (*Poaceae*). Cereals are rich in carbohydrates, proteins, minerals, vitamins, and dietary fibers, which contribute to ∼47% of dietary protein, ∼51% of caloric intake, and ∼5-7% of edible fat in human diet (Gudi et al. 2022; Borrelli and Ficco 2025). Rice, wheat, maize, oat, barley, and sorghum, are the major cereal crops grown globally, among which wheat, oat, and barley require cool environments (Loskutov 2021). Cereal crops are the most abundantly grown and consumed food on the earth with annual production of ∼2850 million metric tons (https://www.fao.org). Despite their economic and nutritional significance, global production of cereal crops is severely affected by various biotic and abiotic stresses. Major biotic stresses include fungal diseases such as rusts, blast, and *Fusarium* head blight as well as insect-pests like hessian fly, shoot fly, brown plant hopper (BPH), wheat stem sawfly, aphids, stem borer, and bacterial diseases such as bacterial leaf streak (BLS) (Chandra et al. 2024; Singh et al. 2024, 2025). Key abiotic stresses affecting cereal crops include drought, heat, salinity, lodging, ion toxicity, etc. (Jeyasri et al. 2021; Tanin et al. 2022; Singh et al. 2023; Gudi et al. 2024a, 2025a-d; Rajamanickam et al. 2024).

Drought stress is characterized by a long spell of water scarcity in crop root zones resulting from the lack of enough precipitation coupled with high evapotranspiration. It affects plants morpho-physiological, biochemical, and molecular processes at various developmental stages including, seed germination and seedling establishment, tillering, vegetative, reproductive, and grain filling stages (Begna et al. 2022; Gudi et al. 2024b). Drought stress affects both the arial and underground parts of the plants. In the arial parts, drought stress reduces leaf water content, which results in the drooping and folding of leaves. It also induces partial or complete closure of stomata, which increases leaf temperature and reduces gas exchange (Lv et al. 2023). High leaf temperature disturbs cellular homeostasis by destabilizing proteins and inactivating key enzymes involved in metabolic processes such as respiration and photosynthesis (Asargew et al. 2024). Under severe drought, the arial parts of plants fail to produce enough photosynthates to support root growth, resulting in the short and thick roots (Siddiqui et al. 2021; Kalra et al. 2024). Disturbed root system architecture impairs nutrient and water uptake, further contributing to stunted plant growth. Prolonged and intense drought stress may cause plant wilting and ultimately causing plant death. The impact of drought stress depends on its duration (short and long duration), intensity (mild and severe drought), crop growth stage as well as the crop species and their genotypes (Begna et al. 2022). For instance, drought stress can reduce wheat yield by up to ∼20%, ∼47%, and ∼65%, during booting, tillering, and reproductive stages, respectively (Nezhadahmadi et al. 2013; Gudi et al. 2024a). Similarly, drought stress can cause yield loss of up to 44% in two-row barley and up to 48% in six-row barley (Berki et al. 2025). In general, drought stress reduces grain yield in cereals by negatively affecting yield related traits, such as plant height, effective tiller numbers, spike/panicle length, spikelet number, floret fertility, grains per spike, grain length, grain width, and grain weight.

Negative effects of drought stress can be overcome by developing drought tolerant cultivars in different cereals. However, the narrow genetic diversity for drought tolerance in modern cultivars pose a significant challenge to incorporate favorable alleles into breeding programs (Maccaferri et al. 2019; Jayakodi et al. 2024). This can be addressed by introducing novel genetic variations from the unexplored genetic resources such as global germplasm collections, traditional landraces, and wild relatives, specifically those collected from the hot and dry climates. These germplasm lines have evolved through natural selections over thousands of years, making them inherently adapted to harsh environments (Zhang et al. 2025). As a result, they serve as a reservoir for rich allelic diversity encompassing several adaptive traits that can be harnessed efficiently to enhance drought tolerance. However, efficient exploration and utilization of this huge genetic diversity require rapid, reliable, and scalable germplasm characterization strategies capable of evaluating hundreds to thousands of germplasm lines in short time. Over the last century, several protocols for drought screening have been suggested and utilized in different cereals. These include inducing osmotic stress using polyethylene glycol (PEG-6000) or mannitol, detached plants or leaf assays, greenhouse-based seedling survival assays, utilizing movable rainout shelters, and performing field trails under natural drought conditions (Clawson et al. 1986; Seki et al. 2002; Longenberger et al. 2006; Możdżeń et al. 2015; Gudi et al. 2024a, b; Sapkota et al. 2025). Each of these protocols have their own limitations. For instance, PEG and mannitol induce drought stress by reducing water potential, thereby imposing an osmotic stress, which makes it difficult to interpret the drought stress mechanism (Töpfer et al. 2024). Additionally, these solutes fail to mimic the real-world drought stress conditions. Hence, field-based drought experiments conducted under natural conditions represents the most realistic and effective way of drought screening (Bidinger 2002; Pantha et al. 2025). However, environmental variabilities imposed under field trials such as unpredictable rainfall and varying degree of soil moisture fail to control uniform drought stress throughout the crop cycle and over the years. Under such varying environmental conditions, germplasm characterization for quantitative traits is very difficult, as they are strongly influenced by genotype-environment interaction (Eltaher et al. 2021). Moving rainout shelters offers a partial solution by closing shelters during rainy days and thereby impose controlled drought conditions. However, these shelters are limited in scalability, occupy a smaller number of germplasm lines and incur huge investment costs, which is not affordable by all researchers (Clawson et al. 1986). Most of these methods focus on measuring quantitative traits such as shoot height, root length, tiller number, biomass, grain number, and grain yield. While these traits are agronomically important, they are labor-intensive and time consuming to measure, and subject to high variability. Simplifying and scaling up phenotyping by discovering qualitative traits accelerate large germplasm collections screening as well as identification and characterization of drought tolerance genes. Despite decades of research, there are no standard protocol exists for drought screening across cereal crop species. Considering the impact of drought stress and maintaining the global food security, it is important to develop a robust, cost-effective, and reproducible drought screening protocols suitable for multiple cereals.

To accomplish this, we developed a rapid and cost-effective protocol for screening multiple cereal crops under drought stress at the seedling stage. Using *in-house* developed method, we evaluated 1200 germplasm lines comprising diversity panels of wheat, oat, and barley and the wild relatives of wheat such as *Aegilops tauchsii* and *Ae. umbellulata*. This protocol facilitates efficient and high-throughput evaluation of large number of germplasm lines across different cereals within a short period of time, may thereby accelerate the development of drought resilient cereals for drought-prone environments.

## 2. Materials and methods

### 2.1 Plant materials

In this study, we evaluated a total of 1200 germplasm lines belonging to diversity panels of wheat (419 genotypes), oat (273 genotypes), barley (197 genotypes), and the wild relatives of wheat including *Ae. tauschii* (290 genotypes) and *Ae. umbellulata* (21 genotypes). In wheat, we included both tetraploid (19) and hexaploid (400) accessions. Tetraploid wheat lines comprises genetically and phenotypically diverse accessions belonging to *T. turgidum* ssp*. durum* (12 accessions), *T. turgidum* ssp*. dicoccum* (2 accessions), *T. turgidum* ssp*. dicoccoides* (3 accessions), and *T. turgidum* ssp*. carthlicum* (2 accessions) (Gudi et al. 2026). Hexaploid wheat germplasm lines represent a subset of hard red spring wheat (HRSW) diversity panel (400 accessions) representing cultivars, landraces, and breeding lines (Szabo-Hever et al. 2025). Oat germplasm lines include the subset of global assembly of landraces belonging to “Oat Landrace Diversity (OLD)” panel and the wild and domesticated oats previously used in pangenome and pantranscriptome studies (Rahman et al. 2025; Avni et al. 2026). The barley germplasm lines were drawn from the world barley core collection (BCC), representing accessions collected from different geographical regions (Karki et al. 2024). Similarly, *Ae. tauschii* and *Ae. umbellulata* accessions represents the global collections of wild wheat relatives (Cavalet-Giorsa et al. 2024; Singh et al. 2024).

### 2.2 Experimental design and drought screening

We standardized a protocol for screening multiple cereal crops under drought stress. The detailed procedure of *In-house* developed drought protocol is presented in Figure 1. In this protocol, we used experimental pots measuring 10.5cm L × 10cm W × 12cm H. Each pot was tared to zero before being filled with 200g of soil (PRO-Mix BRK20 Biofungicide + Mycorrhizae peat/bark mix) and were watered to full saturation to maintain uniform moisture content in all pots. Healthy and bold seeds from each genotype were sown in three pots representing three replications. Usually, 13 seeds were sown in each pot for small and round seed species (such as wheat), and 8 seeds were sown for long and large seed species (such as oat and wild relatives of wheat). Once sowing is completed, pots were arranged in randomized complete block design (RCBD). Throughout the experiment, pots were placed on mesh-trays (10 pots in each tray) supported by water holding trays below. To minimize the environmental effects resulting from exposure of edges of trays or benches to light, airflow and temperature, a border row was planted using pots of identical size (Supplementary Figures 1 and 2). Once seeds were germinated (usually 3-4 days after sowing in cultivated species and 5-6 days in wild species), extra seedlings were thinned by maintaining uniform number of seedlings in each pot. Precautions were taken to retain the uniform and healthy seedlings by removing defected and late germinated seedlings. Total 10 seedlings were maintained in each pot for small and round seed species, while five seedlings were retained for long and large seed species. After thinning, 5g of slow-release fertilizer (Osmocote Plus 15-9-12 patterned release fertilizer) was added to each pot. Following fertilizer application, pots were watered by sprinkling from the top. Thereafter, watering was done every two days for cultivars and landraces, and every 3-4 days for wild species. Once plants reached two leaf stage (i.e., ∼10-12 days after sowing in cultivars and landraces and ∼14-15 days in wild relatives), regular watering has been done along with filling water holding trays. Pots were allowed to absorb water for overnight until full saturation. The following day, water holding trays were removed and drought stress was initiated by withholding water until volumetric moisture content of soil reached to five percent and some of the genotypes started to show wilting symptoms. Once drought stress period is completed, pots were rewatered to full saturation and allowed for two days recovery.

**Figure 1.**
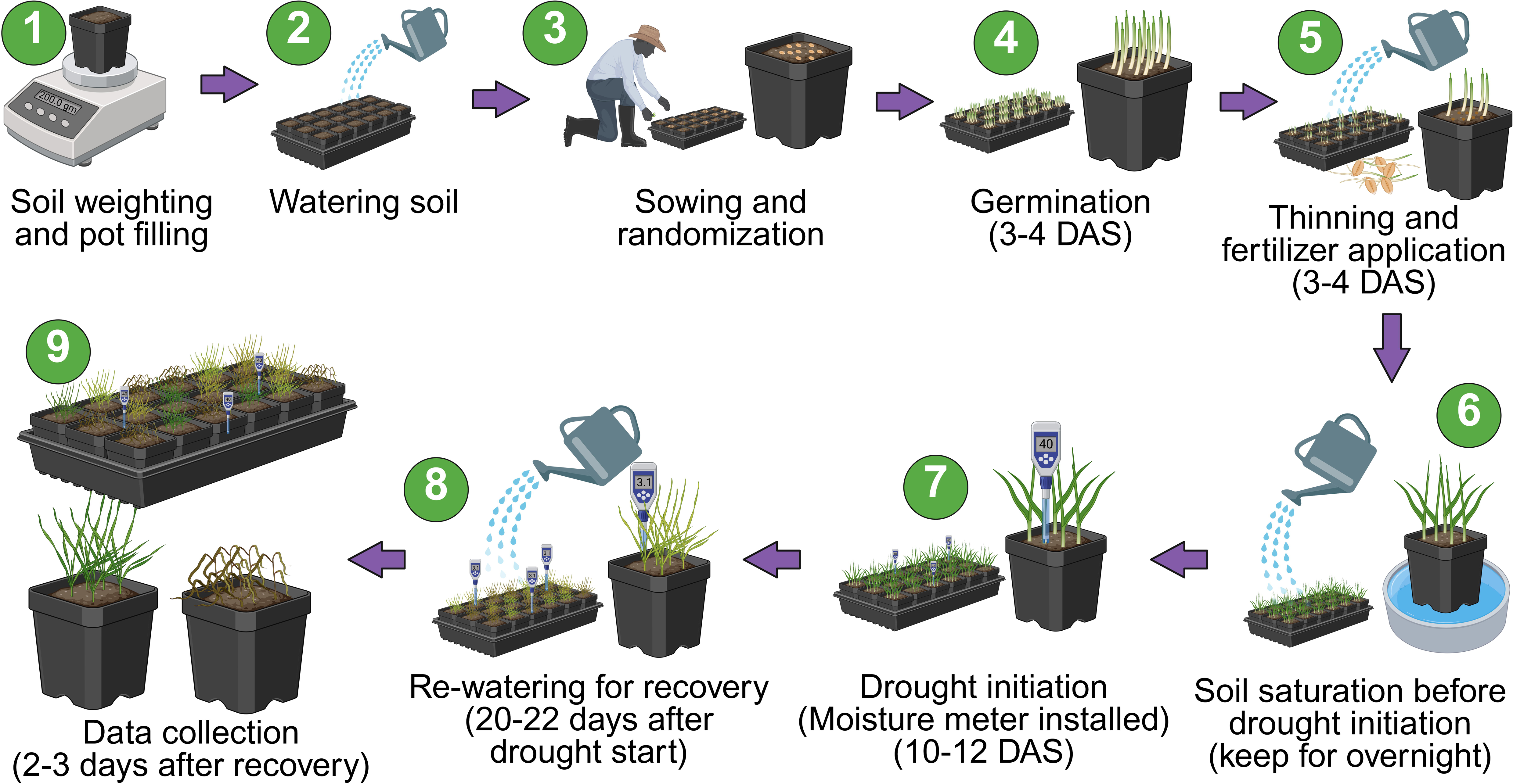
Schematic diagram illustrating the experimental workflow for drought screening at seedling stage. **DAS**, days after sowing

### 2.3 Data collection

The wilting score (WS) was recorded for individual plants in each replication/pot based on the performance of top three leaves using 0-6 scale, suggested in Gudi et al. (2026). “WS 0” means no wilting symptoms with all top three leaves healthy; “WS 1”, the top two leaves healthy with partial wilting of third leaf; “WS 2”; the top two leaves healthy with complete wilting of third leaf; “WS 3”, the top one leaf healthy with partial wilting of second leaf and complete wilting of third leaf; “WS 4”, the top one leaf was healthy with complete wilting of second and third leaves; “WS 5”, partial wilting of the topmost leaf and complete wilting of second and third leaves; and “WS 6”, a completely wilted or dead plant (Figure 2). Genotypes with less than ten or five seedlings per pot or those having one or more weak, chlorotic, stunted or mechanically damaged seedlings were excluded from the study to avoid potential bias in the results.

**Figure 2.**
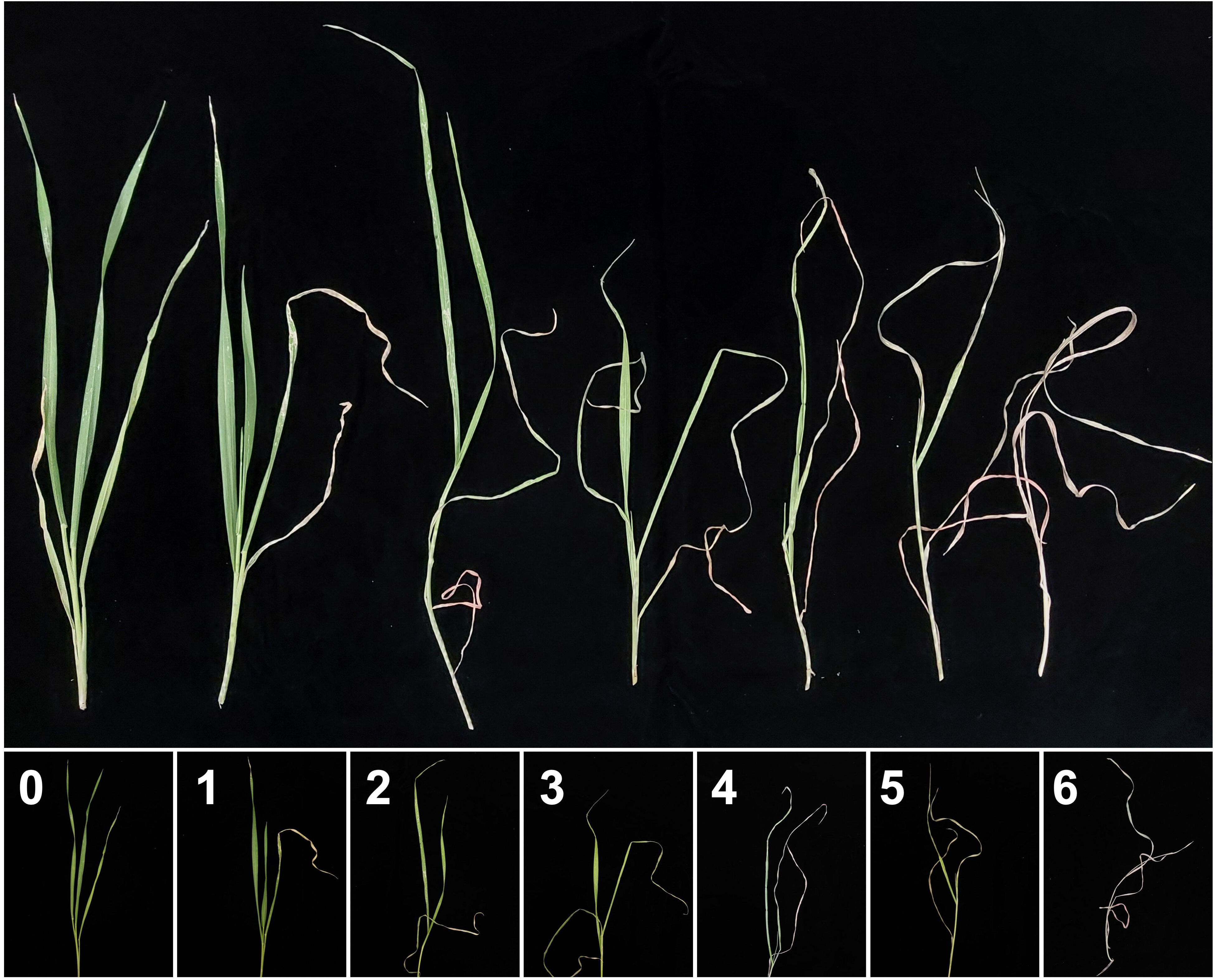
Wilting score (WS) scale used for phenotypic data collection under drought stress. “WS 0” represents extreme drought tolerance and “WS 6” represents extreme drought sensitive.

### 2.4 Greenhouse settings

All experiments were conducted in the USDA-NCSL greenhouse facility at Fargo, North Dakota, USA (46.9002° N latitude, 96.8032° W longitude). Growing conditions inside the greenhouse were set to 16- hour light with 25 °C day temperature and 8-hour darkness with 22 °C night temperature. Relative humidity (RH) was maintained at ∼65%.

### 2.5 Adult plant drought stress screening

Two contrasting hexaploid wheat genotypes identified from seedling stage evaluation-drought tolerant FLD and sensitive J5 genotypes-were evaluated for their response to adult stage drought stress. The experiment was conducted in randomized complete block design (RCBD) with three treatments: (i) well-watered control; (ii) drought stress imposed at anthesis stage; and (iii) drought stress imposed during grain filling stage, each with three biological replicates per genotype. The experimental setup thus comprised of total 18 pots including nine for each genotype (three treatments × three replications). Three bold and healthy seeds from each genotype were sown in six-inch pots with drainage holes in the bottom and seedlings were thinned to one plant per pot to eliminate intra pot competition. Before sowing, the individual pots (15 cm in diameter and 18 cm in height) were tared to zero and filled with 550 g of soil to ensure uniform soil quantity across the pots. All plants were grown under optimal growing condition until anthesis stage by providing regular irrigation and applying 25 g of slow-release fertilizer (Osmocote Plus 15-9-12 patterned release fertilizer).

For anthesis-stage drought treatment, irrigation was withheld from three pots per genotype (three replicates) beginning at the anthesis stage. For the grain filling stage drought treatment, irrigation was withheld similarly from three pots per genotype at the grain filling stage. Volumetric soil moisture content of each pot was regularly measured by installing Zynect Soilmote (www.zynect.com) wireless soil moisture sensors. When soil moisture content of the pots reached 5%, pots were re-watered to field capacity to allow for recovery, after which normal irrigation was resumed for the remaining growth cycle. At maturity, individual plants were harvested, and data was recorded on different agronomic traits including, seeds per spike (SPS), seed weight per spike (SWPS; g), effective tiller number (ETN), and grain yield (GY; g).

### 2.6 Volumetric soil moisture (VSM) content monitoring

Soil moisture content in the pots was tracked throughout the drought period by installing Zynect Soilmote wireless soil moisture sensors. Moisture meters were programmed to record soil water content at 15 minutes interval, generating 96 data points per day. Soil moisture data monitored regularly through “Zynect Sensors” mobile application connected via Wi-Fi. The application enabled real time monitoring of sensors performance with push notifications if any device detected errors or if soil moisture fell below the predefined threshold (<5%) or exceeded the upper threshold (>50%). Upon the completion of experiment, the recorded data were exported via email and downloaded for subsequent analysis.

The mean daily volumetric soil moisture (VSM) content was calculated by averaging the 96 measurements collected over 24 hours period. Then, the relative soil moisture (RSM; in %) content was estimated using the formula:

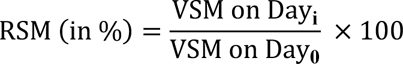

Where, ‘RSM’ is the relative soil moisture content of the pot (in %), ‘VSM on Day_i_’ is the volumetric soil moisture content of the pot on i^th^ day, and ‘VSM on Day_0_’ is the volumetric soil moisture content of the pot on 0^th^ day (i.e., initial soil moisture content).

For estimating the soil drying index (SDI in %), we used:

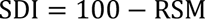

Where, ‘SDI’ is the soil drying index (in %) and ‘RSM’ is the relative soil moisture content of the pot (in %).

The RSM and SDI were plotted using the ggplot2 package of RStudio (Ginestet 2011).

### 2.7 Statistical analysis

Raw WS data were subjected to mixed linear model (MLM) to extract best linear unbiased prediction (BLUP) using LME4 package of RStudio (Kronthaler and Zöllner 2021). BLUP values were calculated using the following model:

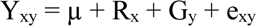

Where, Y_xy_ represents seedling trait of interest, µ represents overall mean, R_x_ represents effect of x^th^ replication, G_y_ represents effect of y^th^ genotype, e_xy_ represents error associated with x^th^ replication and y^th^ genotype, which is assumed to be normally and independently distributed, with mean zero and homogeneity of variances as V.

Broad-sense heritability (h^2^_bs_) was calculated using the following formula:

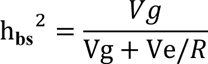

Where, h _bs_^2^ represents broad-sense heritability, Vg represents genetic variance, Ve represents error variance, R represents number of replications.

Coefficient of variation (CV) was measured using:

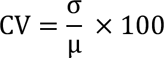

Where, CV represents coefficient of variation (in %), “σ” represents standard deviation and “μ” represents overall mean.

The analysis of variance (ANOVA) based on RCBD design was performed for agronomic traits by considering genotypes, treatments, and their interactions as fixed effects, while replications as block effects. The following model was used:

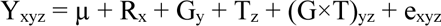

Where, Y_xyz_ represents observed trait value, µ represents overall mean, R_x_ represents effect of x^th^ replication, G_y_ represents effect of y^th^ genotype, T_z_ represents effect of z^th^ treatment, (G×T)_yz_ represents Genotype × Treatment interaction, and e_xyz_ represents random error.

The genotype means were compared using the Fisher’s least significance test (LSD) at the 5% level of significance.

## 3. Results

### 3.1 Assessing the progress of drought stress through relative soil moisture (RSM)

Monitoring relative soil moisture (RSM) content of pots showed similar soil drying pattern across the species (Supplementary Figure 3). At the onset of drought, the SDI was 0% and it increased gradually for initial few days of water withholding, followed by rapid increase during intermediate days until value reaches to 87-95% (Figure 3). Upon rewatering, the SDI declined sharply, indicating full saturation of pots. In addition, the close agreement between individual replicates and mean SDI demonstrates the reproducibility and robustness of protocol across the evaluated species.

**Figure 3.**
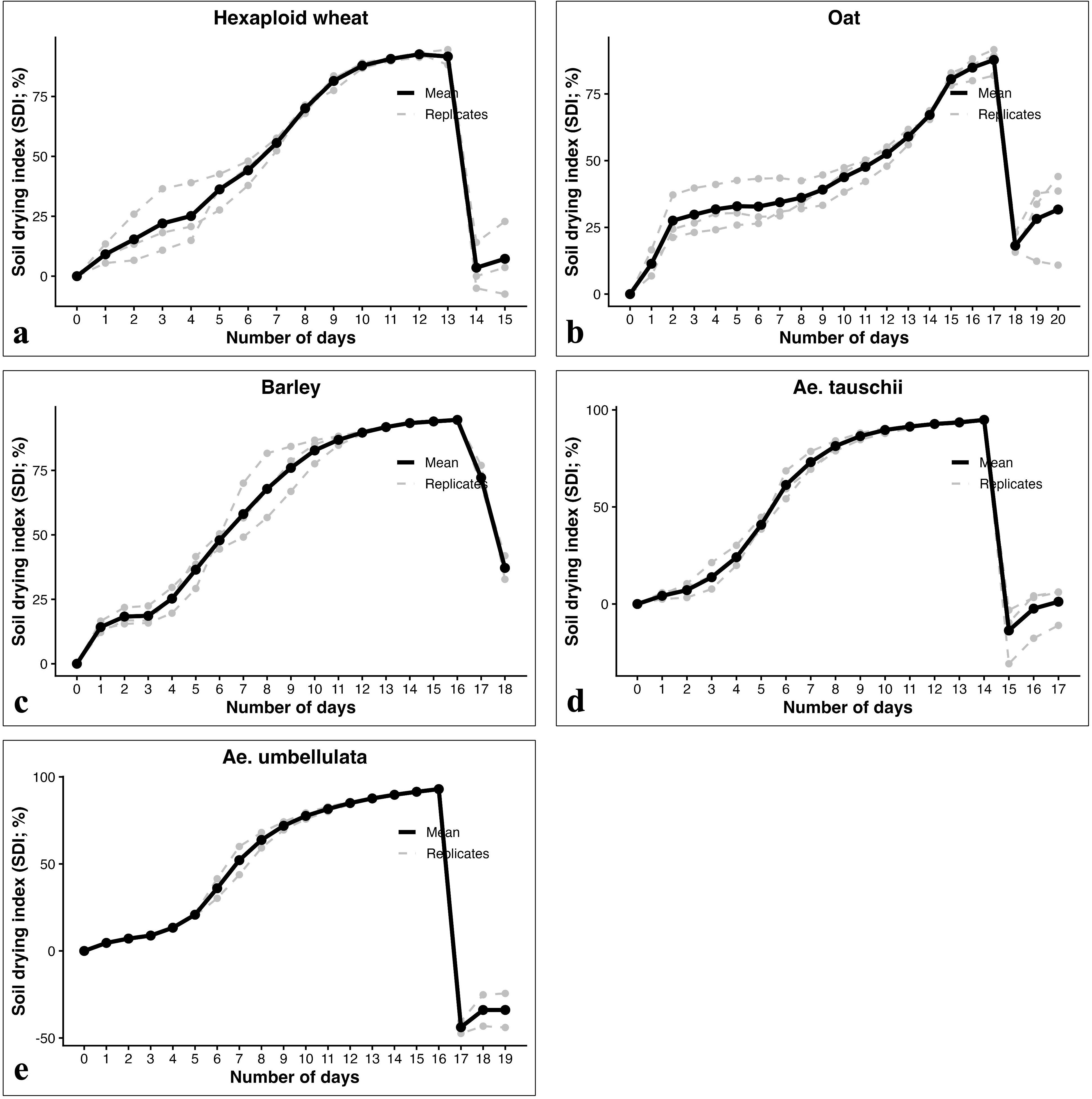
Line graphs showing soil drying index (SDI) of pots during the drought experiment in: (a) wheat, (b) oat, (c) barley, (d) *Aegilops tauschii*, and (e) *Aegilops umbellulata*. Daily SDI represents the mean of 96 measurements recorded at 15 minutes interval. Grey dotted lines indicate the SDI of three biological replicates, and the black dotted lines indicate the mean SDI.

Although overall soil drying pattern was similar among the different species, number of days required to reach maximum SDI varied a little. For instance, oat took highest number of days (17 days) to reach maximum SDI, while wheat took only 13 days to reach maximum SDI (Figure 3). The observed difference in number of days required to reach maximum SDI among different species may be attributed to species-specific differences in water usage, transpiration rate, canopy development, leaf and root architecture.

### 3.2 Phenotypic evaluation for drought stress tolerance

Drought protocol developed in this study produced a clear and reproducible drought symptom across the crop species evaluated. As drying of soil progressed, seedlings exhibited visible drought symptoms including drooping, rolling, yellowing, and wilting of leaves, with symptom severity increasing as SDI approaching maximum value. The initial symptoms were first noticed on old leaves and subsequently progressed to new or young leaves as drought conditions intensified. Although, similar drying pattern was observed across the species, the onset and severity of wilting differed markedly among them. Wheat showed early wilting symptoms, whereas oat seedlings survived for longer period and showed delayed wilting symptoms. Upon rewatering, seedlings showed varying degree of recovery depending on crop species and genotypes.

Analysis of WS data revealed a substantial phenotypic variation among the genotypes within each crop species (Table 1). The CV ranged from 17.93% in *Ae. umbellulata* to 44.94% in *Ae. tauschii*, while broad sense heritability (H²_bs_) ranged from 43% in *Ae. tauschii* to 94.1% in *Ae. umbellulata*, showing differing contribution of genetic and environmental factors to wilting response across the species. Based on mean WS across biological replicates, we classified genotypes into tolerant (WS < 3), moderately tolerant (WS = 3-4), and sensitive (WS > 4). Using these criteria, we identified 3, 185, 50, 56, 74, and 5 drought tolerant genotypes in tetraploid wheat, hexaploid wheat, oat, barley, *Ae. tauschii*, and *Ae. umbellulata*, respectively (Figure 4). Images showing contrasting response of some of the representative drought-tolerant and sensitive genotypes of various species are presented in Figure 5. Together, these results demonstrate the effectiveness of developed protocol in discriminating contrasting drought responsive genotypes across the diverse small grain cereal crops and their wild relatives.

**Table 1.** Descriptive statistical analysis for wilting score (WS) in various populations evaluated in this study.

| Genotypes/population | Number of genotypes | F-value | Pr(>F) | CV (%) | Broad-sense heritability ($H^2_{bs}$) |
| --- | --- | --- | --- | --- | --- |
| Tetraploid wheat | 19 | 7.457 | 1.3E-06 *** | 20.59 | 89.7 |
| Hexaploid wheat | 400 | 3.88 | 2.2e-16 *** | 36.17 | 76.4 |
| Oat | 273 | 4.008 | 2.2e-16 *** | 21.35 | 75.4 |
| Barley | 197 | 2.143 | 1.3e-09 *** | 36 | 56.7 |
| *Aegilops tauschii* | 290 | 1.72 | 2.3e-07 *** | 44.94 | 43 |
| *Aegilops umbellulata* | 21 | 15.005 | 1.045e-11 *** | 17.93 | 94.1 |

CV, coefficient of variance. Significance levels: $P \leq 0.05$ (\*), $P \leq 0.01$ (\*\*), and $P \leq 0.001$ (\*\*\*).

**Figure 4.**
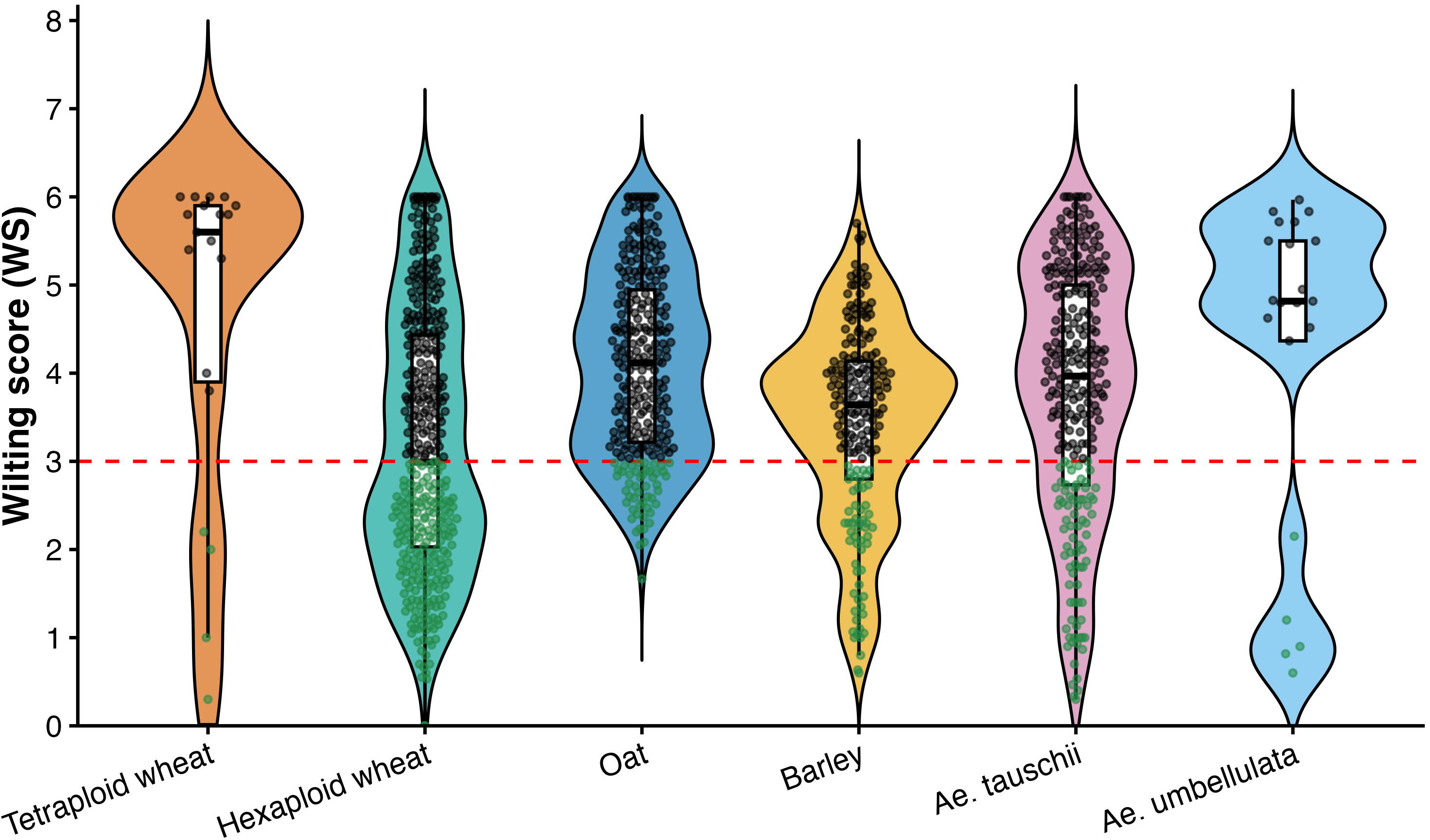
Violin plots showing the distribution of drought response among the diverse genotypes of tetraploid wheat, hexaploid wheat, oat, barley, *Aegilops tauschii*, and *Ae. umbellulata*. Small dots within the violin plots represents individual genotype. The red dotted line indicates the wilting score (WS) threshold used to classify genotypes as drought tolerant and drought sensitive. Genotypes with WS < 3, shown as green dots, were classified as drought tolerant.

**Figure 5.**
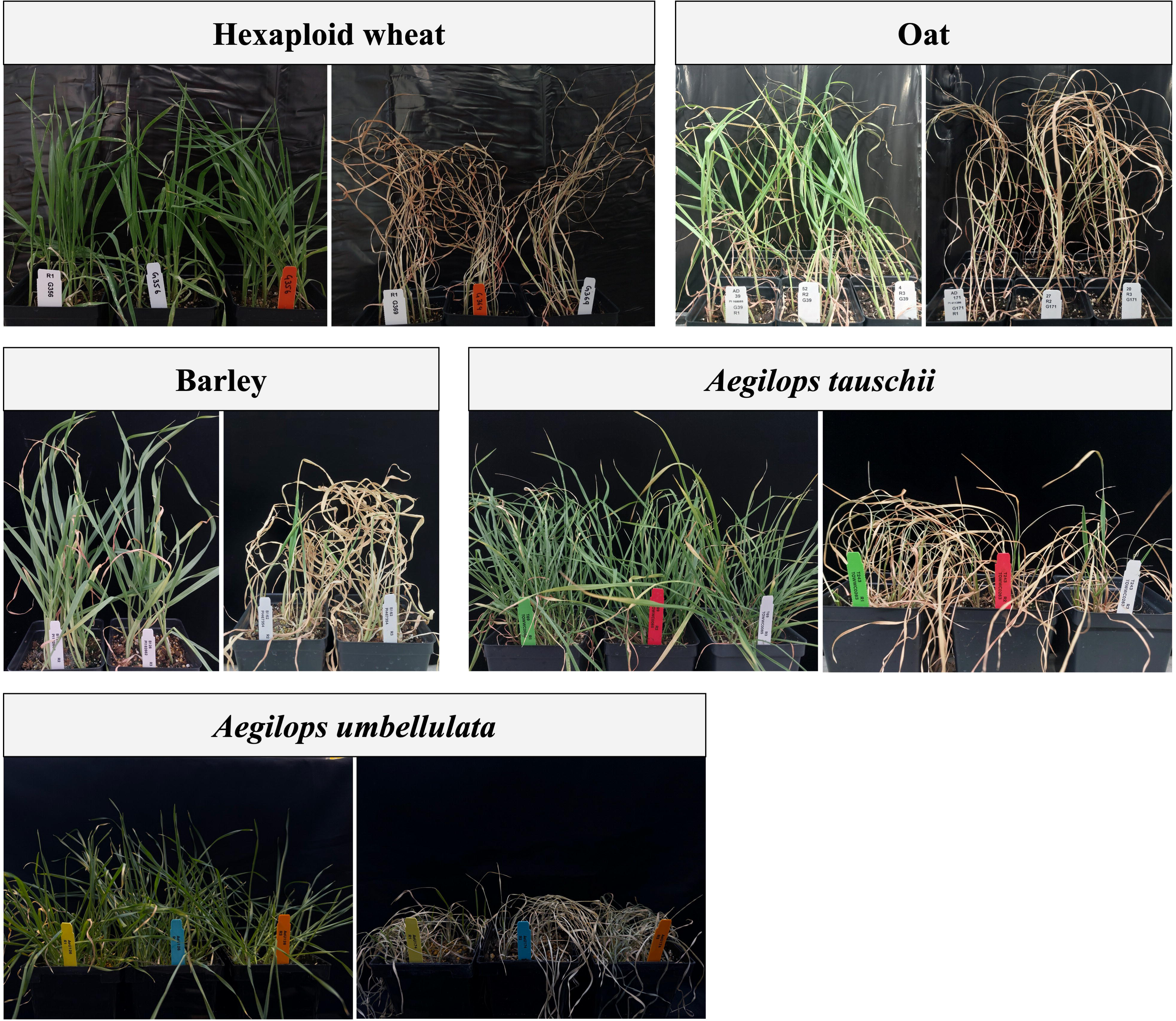
Representative images of extreme drought tolerant (left) and drought sensitive (right) genotypes after drought treatment in wheat, oat, barley, *Aegilops tauschii*, and *Ae. umbellulata*.

### 3.3 Reproducibility of the drought protocol

To assess the reproducibility of the drought protocol developed in this study, a subset of contrasting genotypes identified from the diversity panels were re-evaluated in an independent experiment using the same procedure. For this purpose, we selected three drought tolerant (PI 478742, PI 481521, and Altar 84) and one drought sensitive genotypes (Langdon 16) of tetraploid wheat, one drought tolerant (FLD) and drought sensitive (J5) genotypes of hexaploid wheat, and one drought tolerant (PI 168088) and drought sensitive (CIav 9101) genotypes of oat.

Drought responses observed in the preliminary evaluation were consistently reproduced in the repeated experiment (Figure 6). Drought tolerant genotypes recovered very well after rewatering with low WS (WS < 3), while drought sensitive genotypes died completely and displayed high WS (WS > 5). Differences in the mean WS between initial screening and repeated experiment were very low, with drought tolerant lines displaying WS of 2 and 1.5 for Altar 84, 0.3 and 1.1 for PI 478742, 2.2 and 1.3 for PI 478742, 1.18 and 0.73 for FLD, and 2.33 and 2.31 for PI 168088, respectively. In contrast, drought sensitive genotypes consistently displayed high WS of 5.3 and 5.5 for Langdon 16, 6 and 6 for J5, and 6 and 6 for CIav 9101, respectively, in initial screening and repeated experiment. This consistent separation of tolerant and sensitive genotypes across the independent experiments suggests that the developed protocol is reproducible and provide reliable approach for characterizing global germplasm collections in a relatively short time.

**Figure 6.**
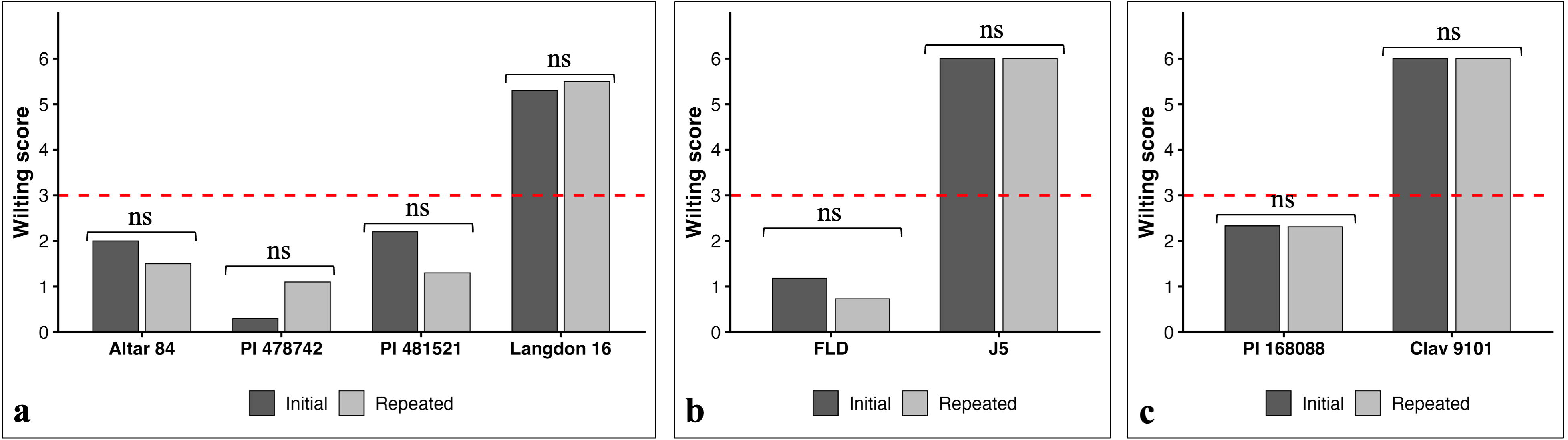
Assessment of the reproducibility of the proposed drought screening protocol using contrasting genotypes of: (a) tetraploid wheat, (b) hexaploid wheat, and (c) oat in independent drought experiment. The genotype means were compared using the Fisher’s least significance test (LSD) at the 5% level of significance. **ns**, non-significant.

### 3.4 Validation of drought protocol at adult stage

To examine whether the contrasting drought responsive genotypes identified in the seedling drought screening maintain similar response at the later developmental stages, we evaluated one drought tolerant (FLD) and one drought sensitive (J5) hexaploid wheat genotypes under drought stress imposed during anthesis and during grain-filling stages (Table 2). The drought tolerant genotype FLD outperformed the drought sensitive genotype J5 both at anthesis and grain-filling stages (Figure 7). Drought stress significantly reduced SWPS (52.47 % and 33.63 %), ETN (38.1 % and 55.04 %), and GY (73.02 % and 19.05 %) during anthesis and grain-filling stages in sensitive genotype J5 (Supplementary Figure 4). In contrast, FLD showed comparatively stable performance, with significant reduction observed only in SWPS (38.24 %) and GY (58.73 %) during anthesis stage drought stress, while the performance of most traits unchanged during grain-filling stage. These results provide evidence that drought protocol developed for seedling stage evaluation identifies contrasting drought responsive genotypes, which also maintain their contrasting performance in later developmental stages.

**Figure 7.**
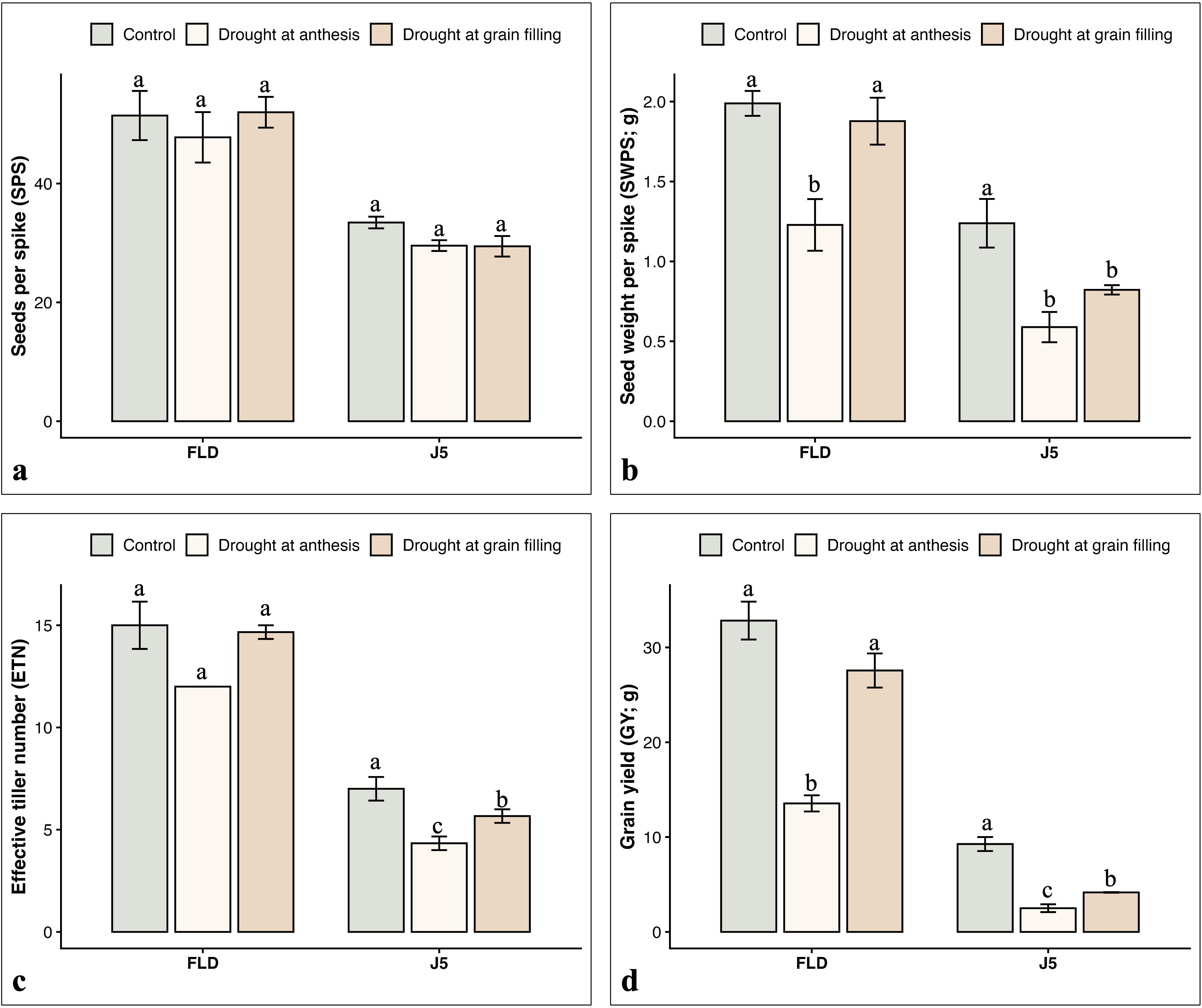
Performances of drought tolerant (FLD) and sensitive (J5) genotypes of hexaploid wheat under control (light green bar) and adult plant drought stress imposed at the anthesis (light yellow bar) and grain filling (light brown bar) stages for: (a) seeds per spike (SPS), (b) seeds weight per spike (SWPS; g), (c) effective tiller number (ETN), and (d) grain yield (GY; g). Different letters (a, b, and c) above the graph represents significant differences among genotype means using the Fisher’s least significance test (LSD) at the 5% level of significance.

**Table 2.** Analysis of variance (ANOVA) for agronomic traits evaluated under control conditions, drought stress imposed at anthesis, and drought stress imposed during grain-filling stage in drought tolerant (FLD) and sensitive (J5) genotypes.

| Trait | Replication | Genotype | Treatment | Genotype:Treatment |
| --- | --- | --- | --- | --- |
| SPS | 0.77 | 1.1e-05 *** | 0.47 | 0.7 |
| SWPS | 0.046 * | 8.66e-07 *** | 0.0002 *** | 0.12 |
| GY | 0.6 | 9.34e-09 *** | 2.41e-05 *** | 0.001 ** |
| ETN | 0.99 | 8.6e-08 *** | 0.011 * | 0.65 |

SPS, seeds per spike; SWPS, seed weight per spike (g); ETN, effective tiller number; GY, grain yield (g). Significance levels: $P \leq 0.05$ (\*), $P \leq 0.01$ (\*\*), and $P \leq 0.001$ (\*\*\*).

## 4. Discussion

Drought is a major environmental stress that limits growth, development, and productivity of cereal crops worldwide (Zhu et al. 2026). Alleviating negative effects of drought stress requires identification of drought tolerant genotypes by developing an efficient screening method capable of evaluating large germplasm collections within short time. Although several drought screening methods have been developed, no single protocol works efficiently across different crop species (Ahady et al. 2025). Moreover, these methods not only require specialized instrumentation, but also are labor intensive and difficult to scale up for evaluating large-germplasm collections in a short time. For instance, field-based screening provides the realistic drought stress conditions, but suffers from uncontrolled spatial and temporal environmental variabilities, often making it difficult to achieve reproducible results (Bidinger 2002). More reproducible methods involve the germplasm evaluation in medium whose water potential is lowered by using osmotic agents such as poly-ethylene glycol (PEG) or mannitol (Gudi et al. 2024a, b). Such method is suitable for studying short-term drought responses only, as studying long-term drought stress is affected by phytotoxicity of osmotic agents with difficulty in maintaining constant osmotic potential throughout the experiment (Verslues et al. 1998). Therefore, developing a rapid, efficient, and reproducible soil-based drought screening protocol that balances environmental control with biological relevance is essential.

In this study, we optimized a high throughput, efficient, and highly reproducible drought protocol for screening hundreds to thousands of germplasm lines of small grain cereals and their wild relatives at seedling stage. Maintaining uniform environmental conditions throughout the experiment is essential for achieving reproducible results (Moshelion et al. 2024). Here, continuous monitoring of soil moisture content revealed a similar soil drying pattern across the biological replicates suggesting that the environmental variations during the experiment were minimal. Although minor differences in drying period were observed among the species, these disparities likely reflect the inherent variation in stomatal regulations, canopy development, leaf and root architecture of different species, rather than environmental variations (Pataki and Oren 2003; Wright et al. 2024). The use of uniform sized seeds, equal quantity of soil and fertilizer, an equal number of healthy seedlings, controlled watering, and border rows, all contributed to the reduced environmental variations.

Wilting represents the cumulative effect of leaf water status, reduced cell turgor, and impaired cellular integrity, metabolic activity and physiological processes under drought stress (Zhang et al. 2024). It integrates multiple physiological responses into single and easily measurable trait, as wilting symptoms developed gradually with proportional decline in soil moisture content (Vennam et al. 2026). Moreover, visual scoring based on leaf or whole-plant wilting enables rapid evaluation of several germplasm lines without requiring specialized expertise. To exploit this easily measurable trait, we developed the WS scale ranging from 0-6, as suggested in Gudi et al. (2026), based on the performance of top three leaves under drought stress. Phenotypic evaluation using this scale revealed a substantial genetic variation among the germplasm lines within each species and identified contrasting drought tolerant and sensitive genotypes, which demonstrates the ability of the developed protocol in discriminating contrasting genotypes.

Achieving repeatability in the performance of genotypes across independent experiments is critical for selecting drought tolerant genotypes for inclusion in the breeding program with high confidence (Moshelion et al. 2024). Reevaluation of representative extreme genotypes of tetraploid wheat, hexaploid wheat, and oat revealed consistent drought response in an independent experiment, indicating high reproducibility of the developed protocol.

The relationship between seedling stage and adult plant drought tolerance remains a subject of ongoing debate, as genotypes exhibiting drought tolerance at the seedling stage may not always show similar response at the later developmental stages (Harrison and LaForgia 2019). Consequently, introducing a seedling stage drought tolerant genotype into breeding program may confer tolerance to early-stage drought stress only, without providing protection against drought occurring during later developmental stages. Therefore, it is essential to identify the genotypes with stable performance across the developmental stages, and developing a protocol capable of identifying seedling stage tolerance that persist into later developmental stages is of particular importance. To assess the efficacy of our protocol in this regard, the contrasting genotypes identified at seedling stage were further evaluated under drought stress imposed during anthesis and grain filling. The drought tolerant genotype consistently outperformed sensitive genotype at both stages, demonstrating the contrasting genotypes identified by our protocol discriminated against not only at seedling stage, but also at later, agronomically critical stages of development. These results provide evidence that the developed protocol serve as an efficient preliminary screening method for identifying contrasting drought responsive genotypes for early and later developmental stages.

### 4.1 Potential applications of the proposed protocol

The proposed drought protocol provides a rapid and reliable approach for preliminary evaluation of diverse germplasm lines of small grain cereals and their wild relatives. Owing to its simplicity and high-throughput nature, this protocol enables the evaluation of hundreds to thousands of accessions within a short time, making it well suitable for germplasm characterization in breeding programs. Since, experiments are conducted under controlled greenhouse facilities, environmental influences are very less, resulting in high reproducibility. Furthermore, scoring plants using a WS scale simplifies the phenotyping step, thereby reducing the need for labor-intensive measurements or specialized equipment. Additionally, by classifying diverse germplasm into tolerant, moderately tolerant, and sensitive groups, our protocol provides the practical farmwork for downstream genetic studies, such as quantitative trait loci (QTL) mapping, genome-wide association studies (GWAS), and candidate gene analysis.

### 4.2 Drawbacks of the protocol

Although this protocol provides an efficient and reproducible approach for evaluating different cereal crops under seedling drought stress, the major limitations must be addressed. First, conducting large-scale drought experiment is labor-intensive as it requires standard soil preparation, accurate filling of thousands of pots, sowing equal number of seeds in each pot, and thinning excess seedlings. Second, this protocol is designed for seedling stage evaluation only, therefore, the stable performance of genotypes must be confirmed by evaluating the identified genotypes under drought stress at the later developmental stages and at the field conditions. Third, since experiments are conducted under controlled greenhouse facilities, imposed stress doesn’t mimic the real complexities and variabilities of drought stress under field conditions. Finally, the protocol primarily focuses on collecting visual data on the WS, which enables rapid evaluation of thousands of germplasm lines in short time, but it does not assess the underlying morphological, physiological, and biochemical mechanisms of drought tolerance. However, this issue can be overcome by conducting detailed morpho-physiological and biochemical analysis on the selected contrasting germplasm lines.

### 4.3 Precautionary measures for successful implementation of the protocol

Successful implementation of proposed protocol requires careful attention to several factors. First, each pot should be filled with equal amount of soil, as differences in soil volume alters the water holding capacity of pots. This results in unequal drying rates across the replications and introduces a confounding effect that bias genotype selection. Similarly, before filling the pots, soil should be mixed thoroughly to ensure uniform moisture content, and should be free of stones, sticks, and other debris that may disturb soil water distribution. Second, bold, healthy, and uniform sized seeds from each genotype should be used for planting to avoid variation arising from seed quality and vigor. Following germination, equal number of healthy seedlings should be maintained across all pots, as pots with fewer seedlings dry slowly and may exhibit fake tolerance, whereas pots with more seedlings dry faster and may appear more sensitive. Extra care should be taken during thinning to avoid disturbing root system of the remaining seedlings. Seedlings should be removed completely rather than breaking at the soil surface, as partially removed seedlings may regrow and interfere with genotype true response. Finally, while working with wilds, species-specific germination characteristics must be considered. For instance, a single spikelet of *Ae. tauschii* can produce multiple seedlings arising from the florets of different sizes, and in that case, careful thinning is required to remove seedling originated from smaller floret by keeping those originating from larger florets.

## 5. Conclusion

In this study, we proposed a rapid screening protocol for assessing the seedling response to drought stress using a simple visual WS. Reducing environmental variation by ensuring homogeneity in the experiment is key for achieving high reproducibility across the independent experiments. This protocol works across both cultivated species and wild relatives, suggesting that it can be used to characterize drought response across diverse genetic backgrounds. Adult-stage evaluation further demonstrates the capability of this protocol to discriminate genotypes at different developmental stages. Although our protocol cannot completely replace the adult plant and field evaluation, it can be used for preliminary evaluation of thousands of germplasm lines, with the shortlisted lines can be subsequently tested in field conditions. In conclusion, this protocol offers practical solution for narrowing down large germplasm collections to manageable number of contrasting genotypes, thereby enabling detailed biochemical, physiological, and genetic studies, as well as their introduction into breeding program.

## Supplementary Figures

**Supplementary Figure 1**. Schematic representation of the randomized complete block design (RCBD) used for drought stress screening in the present study. G1-G9 represents the experimental lines and ‘B’ represents the border rows/plants.

**Supplementary Figure 2.** Progression of drought stress during the experiment, showing the different stages of drought response.

**Supplementary Figure 3.** Line graphs showing the relative soil moisture (RSM) of pots during the drought experiment in (a) wheat, (b) oat, (c) barley, (d) *Aegilops tauschii*, and (e) *Aegilops umbellulata*. Daily RSM represents the mean of 96 measurements recorded at 15 minutes interval. Grey dotted lines indicate the RSM of three biological replicates, and the black dotted lines indicate the mean RSM.

**Supplementary Figure 4.** Percent reduction of drought tolerant (FLD) and drought sensitive (J5) genotypes for (a) seeds per spike (SPS), (b) seeds weight per spike (SWPS), (c) effective tiller number (ETN), and (d) grain yield (GY), evaluated under adult plant drought stress.

## Declarations

## Acknowledgments

The authors acknowledge the Cereal Crops Research Improvement Unit, Edward T. Schafer Agricultural Research Center, USDA-ARS, Fargo, ND, USA for providing funding support and greenhouse facilities.

## Availability of data and material

Not applicable

## Code availability

Not applicable

## Author’s contribution (SG, JS, UG, and RG)

Conceptualization, SG and RG; Methodology and protocol standardization, SG; Experimental design, SG; Investigation and data collection, SG and JS; Data analysis, SG; Writing-original draft preparation, SG and JS; Writing-review and editing, SG, UG, and RG; Supervision, RG. All authors have read and agreed to this version of the manuscript.

## Ethics approval and consent to participate

Not applicable.

## Consent to participate

Not applicable.

## Consent for publication

Not applicable

